# Division of labor and low temperatures predict geographic variation in thermal tolerance of a North American paper wasp (*Mischocyttarus mexicanus cubicola*)

**DOI:** 10.64898/2026.09.01.748716

**Authors:** Kristin M. Robinson, Floria M.K. Uy, Kaitlin M. Baudier

**Affiliations:** School of Biological, Environmental and Earth Sciences, the University of Southern Mississippi, Hattiesburg, MS; Department of Biology, University of Rochester, Rochester, NY

**Keywords:** biogeography, climate, division of labor, macrophysiology, Mexican paper wasp, phenology, thermal tolerance

## Abstract

The biogeographic study of organismal thermal performance is fundamental to our understanding of how climate drives evolution. However, despite highly social insects being popular models for such studies, biogeographic comparisons of thermal functional traits seldom consider division of labor. Individuals within cooperative societies can operate in different microclimates, exposing different task groups to different selection pressures. Here we present a study of how thermal tolerance limits in a social paper wasp vary broadly across temperate and subtropical climates in eastern North America, testing whether division of labor between foundresses and non-reproductive workers generates adaptive variation in thermal performance within the colony. Cold tolerance rather than heat tolerance varied more predictably with environmental temperatures across latitudes, with temperate populations experiencing bouts of winter cold coma, unlike in subtropical populations. Within colonies, reproductive foundresses were also more cold-tolerant than workers, enabling them to remain mobile at cooler periods of early spring during crucial tasks of nest construction, before workers emerge. Together, these results broaden our understanding of how climate shapes thermal performance on both biogeographic and social scales.

## Introduction

Climate-change associated shifts in temperature and rainfall can challenge the persistence of many animals, including social species (Fisher et al., 2019; Menzel & Feldmeyer, 2021). The effects of climate change on social insects are of particular concern because small-bodied poikilotherms are more negatively affected by increasing temperatures (Deutsch et al., 2008). Social insects also perform important ecosystem services such as pollination (Brock et al., 2021; Matias et al., 2017), enhancing decomposition (Del Toro et al., 2012), and control of agricultural pests (Brock et al., 2021). Paper wasps in the family Vespidae in particular are argued to be on-par with bees in terms of the value of total ecosystem services rendered, such as pollination, but also providing some services that bees do not such as consumption of agricultural pests (Brock et al., 2021). This clade also displays a wide variety of social forms (Jia & Makino, 2026; West-Eberhard, 1978), rife with useful model systems for exploring social climate adaptations. However, to accurately study the thermal adaptations of a social species, connection between climate and social strategies must be considered (Schürch et al., 2016).

Studies seeking to elucidate the effects of climate change on natural systems often focus on especially change-sensitive species, those that are likely to be negatively impacted by anthropogenic activity. However, studies of climatic adaptations in change-resilient species are also essential to better understand climate-mediated adaptations that sensitive species lack. Here, we investigate the biogeography of thermal performance in an urban, primitively eusocial acculeate wasp with an expansive native range in North America. *Mischocyttarus* is the most speciose genus among the polistine paper wasps, being comprised of more than 240 neotropical species and several species that are adapted to more thermally seasonal subtropical (±23.4° to ±30.0°) and temperate (> ± 30.0°) latitudes (Corlett, 2013; O’Donnell, 2021; Silveira, 2008). *Mischocyttarus mexicanus cubicola* Richards, is one of the few species in this genus native to temperate zones in North America, notable for nesting on the underside of *Sabal palmetto* fronds, and for their abundance in parks and parking lots within urbanized coastal regions of the southeastern United States (Carlson & Fox, 2021; Nacko et al., 2022). The distribution of *M. mexicanus cubicola* includes several islands in the Caribbean and the mainland coastline of North America from southern Florida northward to North Carolina and westward along the gulf states to as far as Houston (Carpenter et al., 2009; H. R. Hermann & J.-T. Chao, 1984; H. R. Hermann & J. T. Chao, 1984; Nacko et al., 2022; Richards, 1978). Across this expanse, populations differ in phenology, with northern populations becoming inactive over much cooler winter months while southern populations in the subtropics experience mild winters and maintain active colonies year-round (Gunnels, 2007; Gunnels et al., 2008; Mora-Kepfer, 2011).

Within *M. mexicanus cubicola*, the frequency of non-kin cooperation among foundresses varies (Miller et al., 2018; Mora-Kepfer, 2014), with non-kin cooperation being more common in high population density areas (Gunnels et al., 2008), in late summer (Gunnels, 2007), and and at lower latitude (Mora-Kepfer, 2014; Table 1). Thus, by studying thermal performance in this model species, we can perform novel tests of hypotheses about the interplay of climate, thermal performance, and these contrasts in social strategy as selection pressures on disturbance-resiliant thermal adaptation.

**Table 1.** Demographics of colonies collected in this study shown in order of highest to lowest latitude site. Total number of nests sampled per site, average per-nest counts for each level of reproductive caste (foundress, worker, and intermediate), average colony size, and average proportion of the colony composed by foundresses are shown below.

|  | foundresses |  | workers |  | intermediate |  | all female wasps |  | colonies |
| --- | --- | --- | --- | --- | --- | --- | --- | --- | --- |
|  | N | mean/col | N | mean/col | N | mean/col | N | mean /col | N |
| SC/GA | 13 | 1.08 | 20 | 1.67 | 7 | 0.58 | 40 | 3.33 | 12 |
| LA | 19 | 1.36 | 13 | 0.92 | 3 | 0.21 | 35 | 2.5 | 14 |
| N. FL | 28 | 2 | 12 | 0.86 | 4 | 0.28 | 44 | 3.14 | 14 |
| S. FL | 29 | 2.23 | 11 | 0.85 | 0 | 0 | 40 | 3.08 | 13 |
| **All sites** | **89** | **1.67** | **56** | **1.06** | **14** | **0.26** | **159** | **3** | **53** |

The climatic variability hypothesis predicts that low-latitude animals living in relatively stable tropical climates have a narrower set of tolerable environmental temperatures than species inhabiting higher latitudes, where temperature changes more substantially with season (Janzen, 1967). Empirical support for thermal variability as a factor shaping thermal tolerance breadth can be seen in not only in cross-latitude studies (Addo-Bediako et al., 2000; Ghalambor et al., 2006; Sunday et al., 2019), but also across smaller scales such as elevation (Bishop et al., 2016; Gaston & Chown, 1999) and even microhabitat (Baudier et al., 2018). However, accurately studying the thermal limits of highly social insects is complicated by the tendency of some insect societies to have adaptive variation in thermal tolerance ranges among specialized task groups that operate in different thermal environments (Cerda & Retana, 1997; Robinson & Baudier, 2024). How this division of labor in thermal tolerance interplays with macrophysiological predictions at broad geographic scales is not understood. We explore this topic in *M. mexicanus cubicola*, asking how thermal tolerance limits vary with latitude, and how these patterns are affected by seasonal specialization between reproductives and non-reproductives.

In *M. mexicanus cubicola* reproductively active females (henceforth “foundresses”) build nests early in the spring either solitarily or as a group in which one ascends to reproductive dominance and the others become subordinate foragers, with shorter-lived workers emerging and taking over the majority of foraging tasks in warmer months (Gunnels, 2007; Gunnels et al., 2008; Litte, 1977). Therefore, we expected foundressess to tolerate more cold but less heat relative to freshly eclosed workers, and for this effect to be less pronounced in subtropical latitudes where winter months and summer months differ less in temperature. The climatic variability hypothesis predicts that wider thermal tolerance breadths evolve among organisms living in more thermally variable, high-latitude environments (Gaston et al., 2009; Janzen, 1967). We investigated to what extent this selection pressure is mitigated by caste-specific tolerance differences in this species, accounting also for colony demographics, body size, and lipid mass as potential covariates.

## Methods

### Field collections

To control for the effects of seasonality, we collected live colonies of wild *M. mexicanus cubicola* from 16 to 21 of May 2021, in four regions that are representative of the range extent of this subspecies across continental North America (Figure 1). To control for potential effects of the degree of site urbanization, within each region we collected from three different parks associated with urban areas of populations greater than 100,000 (Miami, Palm Bay, New Orleans, and Savannah; Figure 1). The three sites in subtropical southern Florida were Kendall (25.694, −80.375), North Miami (25.846, −80.229), and Davie (26.074, −80.275). The three subtropical northern Florida sites were Fort Pierce (27.412, −80.391), Vero Beach (27.641, −80.455), and Palm Bay (27.998, −80.626). The three southeastern Louisiana sites span the transition point at 30° latitude between subtropical and temperate climates (Corlett, 2013), and included Jean Lafitte Barataria Preserve (29.784, −90.115), Uptown New Orleans (29.934, −90.135), and New Orleans City Park (30.003, −90.094). The three northernmost temperate sites, henceforth referred to as South Carolina-Georgia, were Skidaway, Georgia (31.988, −81.025), Pooler, Georgia (32.110, −81.240), and Hampton Park, South Carolina (32.800, - 79.954).

**Figure 1.**
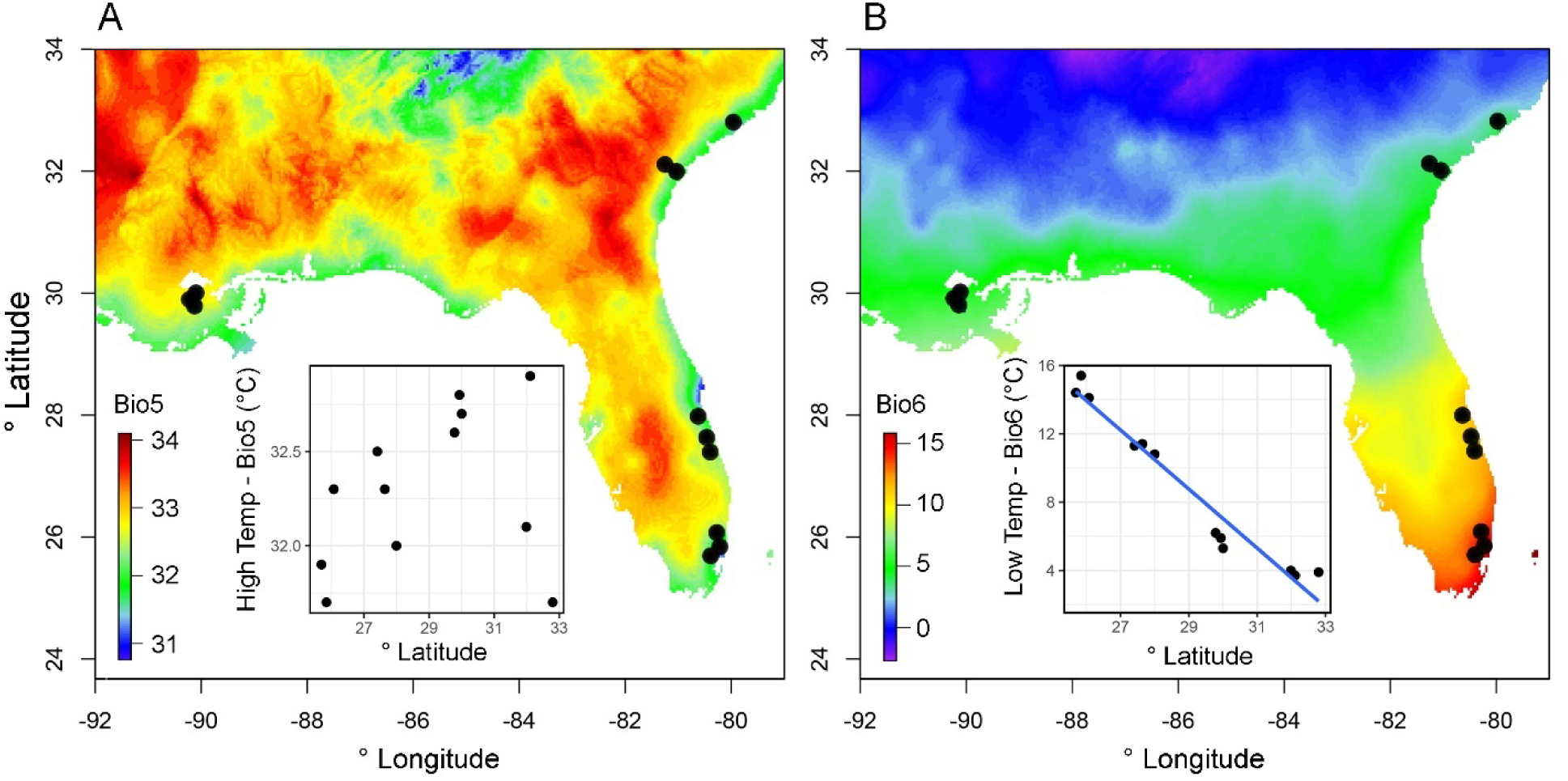
Climate maps of the 12 study sites in 4 regions showing **(A)** latitude as poor predictor of highest temperature of hottest month (Bioclim variable BIO5), and **(B)** the stronger negative correlation between latitude and lowest temperature of coldest month (Bioclim variable BIO6).

Colonies were collected at times associated with low foraging (at or after dusk and/or on rainy days) to maximize the number of captured colony members. All nest materials, brood, and present adult members of each colony were collected into the same nest chamber. Adult males were collected in low numbers and so were excluded from analyses. In this species, the first offspring of early season nests emerge as workers, and there are distinct patterns of cell growth associated with a single versus co- foundress nests (Mora-Kepfer, 2011), attributes we leveraged to identify the castes of collected individuals. Foundresses were identified as females who displayed dominant behavioral traits such as persistent position on the central underside of the nest, as described by Mora-Kepfer (2014), and which had lighter eye color, indicating a more advanced age. Workers were identified as young females (black eye color indicating < 5 days of age), smaller size and/or those that did not display these dominant behaviors. Nest architecture provided corroborated our behavioral and morphological assessment of whether there were single or multiple foundresses in each collected group. Developmental stage of the nest acted as an independent source of caste delineation. A small number of female wasps displayed a mixture of these traits and were categorized as “intermediate” (Table 1); these individuals were excluded from analyses testing for differences between workers and reproductives but were included in the mixed-effect categorical site comparisons of thermal tolerance and heating/cooling tolerance across sites.

Latitude and longitude data for each collected colony were gathered using a handheld GPS unit (Garmin, Ltd. Olathe, KS, USA).

Each nest container consisted of a 15 cm diameter plastic deli-cup with a cotton-stoppered test tube of water, and sucrose in the form of sugar cubes provided *ad libidum* at the base. Each nest was affixed via the pedicel to the lid using lab tape. Immediately following field collection, these chambers were placed into portable coolers that buffered temperatures to between 22° C and 27.25° C (as recorded by iButton thermal loggers, Maxim Integrated^TM^, San Jose, CA, USA). All collected colonies were transported via a climate-controlled car to rearing facilities at the University of Southern Mississippi in Hattiesburg, MS within 5 days of collection.

### Thermal tolerance assays

Once in-lab, nest containers were placed in a common incubator set to 26° C with a 13:11 light:dark cycle and humidity ranging from 76% to 98%. A total of 159 adult wasps (across all sites and colonies) were allowed to acclimate to these conditions for 4 days before being subjected to one of two acute thermal tolerance assays. A total of 76 wasps from 41 colonies were used to estimate maximum critical temperature (CT_max_) assays and 83 wasps from 48 colonies were used to estimate minimum critical temperature (CT_min_). Whenever possible, members from each caste and each colony were assigned to both CT_max_ and CT_min_ assays.

We used dynamic methods to estimate critical temperatures (Lutterschmidt & Hutchison, 1997), with standard modifications used for poikilothermic social Hymenoptera of comparably small size (Baudier et al., 2015; Bujan et al., 2020; Diamond et al., 2012; Oberg et al., 2012; Robinson & Baudier, 2024). Live, acclimated wasps were placed into 1.5 ml Eppendorf tubes that were filled at the top with cotton to prevent thermal refuge in the cap. Subjects of CT_max_ assays were placed in dry heat blocks (Thermal-Lok 4-position dry heat bath, USA Scientific, Ocala, FL, USA), where their temperature was increased 1° C every 10 minutes starting at 30° C. Subjects of CT_min_ assays were placed in cooling blocks (EchoTherm™ IC20, Torrey Pines Scientific, Carlsbad, California, USA), their temperature decreased 1° C every 10 minutes starting at 20° C. In both cases, subject wasps were checked at the end of each 10- minute interval for lack of mobility in response to a light tap. The most extreme temperature at which mobility was observed was considered the wasp’s CT_max_ or CT_min_, respectively. During the first batch of thermal tolerance assays, 3 wasps were kept in tubes but not placed within heating or cooling blocks as controls, to test whether tube conditions (rather than thermal stress) caused mobility loss.

### Climatic variables, Warming tolerance, & Cooling Tolerance

Coordinates gathered via handheld GPS units in the field were used to extract 2.5-minute resolution climate parameters from the online extrapolated climate database Bioclim, which is based on 1960-2021 global weather station data (Booth et al., 2014). The two bioclimatic variables used in this study were “BIO5” (average maximum temperature of the warmest month) and “BIO6” (average minimum temperature of the coldest month) (Figure 1). These variables were used to test whether extreme temperatures were significant selection pressures on thermal tolerance across the North American range of *M. mexicanus cubicola*. These two bioclimatic variables were also used to estimate warming tolerance and cooling tolerance.

Warming tolerance and cooling tolerance are metrics of how closely environmental temperatures are to the exceeding the physiological limits of an organism, and are often discussed in the context of climate change (Anderson et al., 2022). Warming tolerance is defined as the difference between CT_max_ and environmental temperature (Deutsch et al., 2008; Diamond et al., 2012). Similarly, cooling tolerance is the difference between CT_min_ and environmental temperature (Slatyer et al., 2016). To calculate warming tolerance, we subtracted the highest temperature of the hottest month (BIO5; Figure 1A) from the CT_max_ values of wasps. We subtracted CT_min_ from the coldest temperature of the coldest month (BIO6; Figure 1B) to estimate cooling tolerance.

### Measuring potential mechanistic covariates: mass & storage lipids

In addition to testing for social and climatic predictors of thermal tolerance, we conducted secondary explorations of whether these observed patterns were mediated by differences in body size or relative quantity of stored lipids. Following the CT_max_ and CT_min_ assays, subject wasps were immediately frozen (- 20° C). Subject wasps were each dried at 50° C for 3 days before their dry mass was recorded to the nearest 0.001 mg using a microbalance. Preliminary tests showed that 3 days was sufficient to fully dry this species at this temperature. We then conducted a gravimetric diethyl ether assay in order to obtain a measure of the storage lipids within each wasp (Tibbetts et al., 2011) Ostwald et al., 2022; Williams et al., 2011). Diethyl ether extracts non-polar fat stores which we targeted as the primary form of energy storage in diapausing insects and most related to thermal adaptation (Dobush et al., 1985; Hahn & Denlinger, 2011). Dry wasps were placed into filter paper and crushed before being placed into 1-dram vials. Each vial underwent a total of 3 24-hour rounds of extraction. Results from preliminary subsamples showed that this was sufficient to fully extract storage lipids. Each round consisted of filling vials with enough diethyl ether to cover the filter paper and wasp (approximately 4.0 mL). After 24 hours, the solution in the vial was discarded and replaced with fresh diethyl ether until the end of the third round, when we placed the vials on a heating block set to 45° C for 24 hours to allow remaining diethyl ether to evaporate.

Specimens were then dried at 50° C for a minimum of 24 hours before obtaining the post-extraction dry weight of the wasp without the filter paper. We estimated the mass of storage lipids for each wasp as the difference between a wasp’s dry mass before and following the serial extractions. Percentage of storage lipids was calculated from the estimated storage lipid mass relative to the total dry body mass.

### Statistical analyses

All statistical analyses were performed in R version 4.1.1 (R Core Team, 2024). We used linear regressions to explore the relationship between latitude and BIOCLIM estimates for highest environmental temperature (BIO5, average highest temperature of the hottest month) and lowest environmental temperature (BIO6, average lowest temperature of the coldest month). Due to non-parametric demographic data, we used Kruskal-Wallis tests to compare colony size and number of foundresses across the four regions in the study.

We used an analysis of covariance (ANCOVA) to test main and interacting effects of climate (BIO5 & BIO6) and reproductive caste (foundress vs. worker vs. intermediate) on wasp thermal tolerance (CT_max_ & CT_min_). Because sampling took place early in the spring when colonies were first being founded, the average colony size was small (Table 1). This mathematically complicated the use of colony ID as a random factor since replication within each caste and colony was low. To avoid pseudoreplication, the ANCOVA was performed on a dataset consisting of average worker and foundress thermal tolerance for each colony (N = 53 colonies). Separate ANCOVAs were performed for CT_max_ and CT_min_.

To test the significance of low environmental temperature (BIO6) and caste as predictors of wasp size (dry mass) we used a type II analysis on a linear model fitted to the same within-colony caste- averaged dataset as previously described. To resolve an issue of non-normality of residuals, we tested the significance of low environmental temperature (BIO6) and caste as predictors of storage lipid mass we using a type II analysis of a generalized linear model (family = Poisson) fitted to the same within-colony caste-averaged dataset. Whether a wasp’s dry mass or percentage of storage lipid mass were predictive of CT_min_ was tested using a type II analysis of a fitted linear model using the same within colony caste- averaged dataset.

Using the same caste/colony-averaged data, we compared heating tolerance and cooling tolerance of all wasps, including intermediates, across the four regions categorically using two separate ANOVAs. When significant, a post-hoc Tukey HSD was used to test the significance of pairwise differences between the four levels. To generate gradient maps for maximum temperature of warmest month (BIO5) and minimum temperature of the coldest month (BIO6), we integrated bioclimatic data for each collecting site from WorldClim at 0.5 (km^2^) resolution using the *raster*, *rgdal*, *sp, rgeos* and *ggplot* R packages (Bivand et al., 2015; Bivand et al., 2017; Hijmans et al., 2015).

## Results

### Climate & Colony demographics across study sites

Among our 12 study sites in the southeastern United States, latitude was not a significant predictor of high temperature (F_1,10_ = 0.9, R^2^ = 0.011, *p* = 0.372; Figure 1). However, latitude was negatively correlated with low temperature (F_1,10_ = 192.6, R^2^ = 0.946, *p* < 0.001, y = 58.8 - 1.7x; Figure 1). We collected and assayed a total of 159 female wasps from 53 colonies (Table 1). In this spring sampling, both average and median colony size across all regions was 3 wasps. Colony size did not significantly differ across the four regions in this study (Kruskal-Wallis *X*^2^ = 1.07, df = 3, *p* = 0.784; Table 1).

However, number of foundresses per colony significantly increased as latitude decreased across the four regions (Kruskal-Wallis *X*^2^ = 8.2529, df = 3, *p* = 0.041; Table 1).

### Climate & caste as predictors of thermal tolerance

Both BIO6 (low temperature) and caste were significant predictors of CT_min_, and there was no significant interaction between these two factors, indicating no significant difference in caste-specific slopes (Table 2; Figure 2). Results of a post-hoc Tukey HSD test showed that foundresses able to retain mobility at significantly lower temperatures than could workers (*p* = 0.028), with poorly sampled intermediate castes showing no significant difference in cold tolerance from foundresses (*p* = 0.977) or workers (*p* = 0.556). Contrastingly, CT_max_ was not significantly predicted by BIO5 (high temperature) or caste; there was also no significant interaction between these two factors (Table 2; Figure 2).

**Figure 2.**
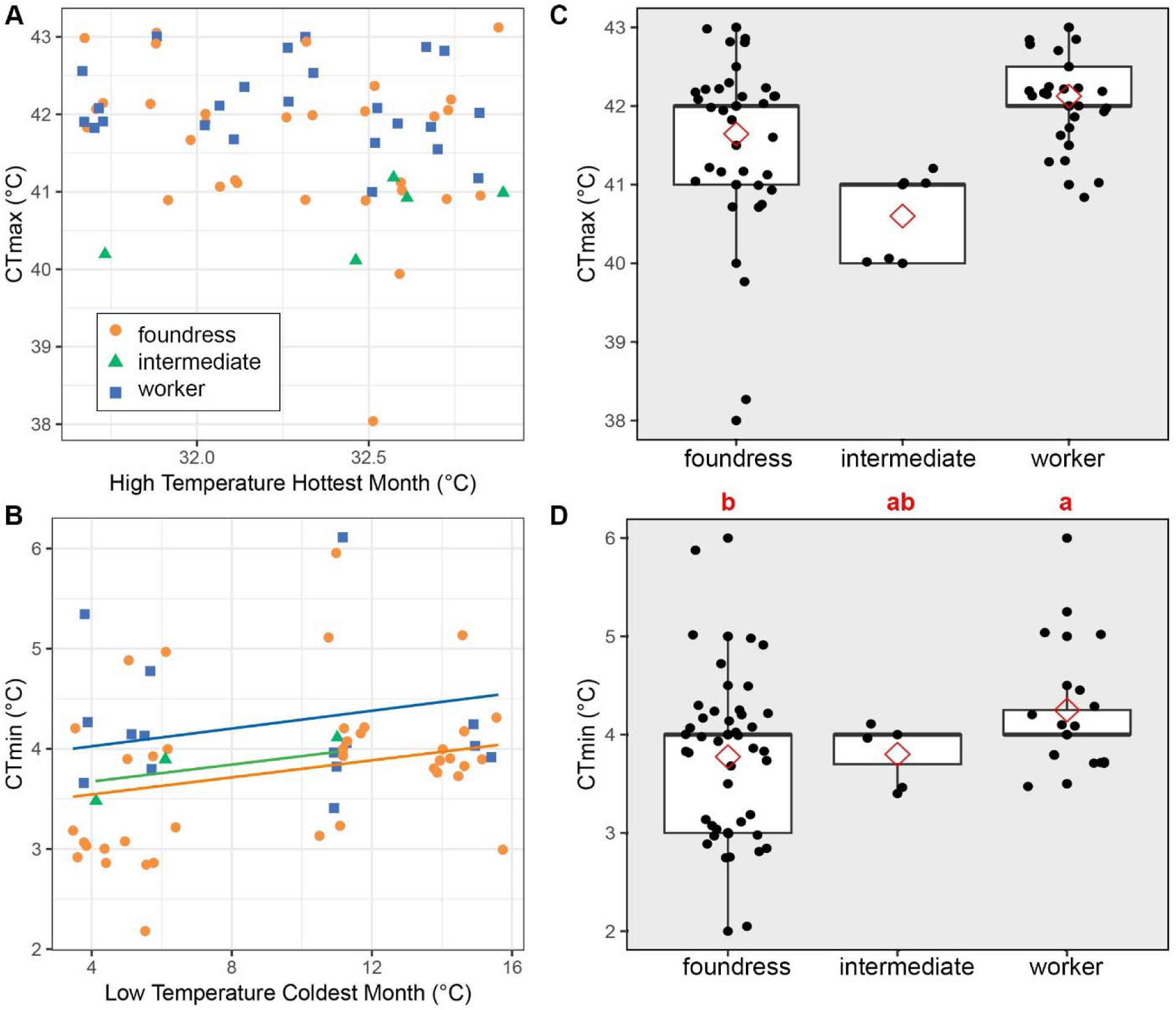
Results of ANCOVA testing the interplay between reproductive caste (foundress vs. worker vs. intermediate) and high temperature (BIO5) or low temperature (BIO6) as predictors of heat tolerance (CT_max_) or cold tolerance (CT_min_), respectively. Statistical outputs of the analysis are shown in Table 2. **(A-B)** Regression lines are shown only for slopes significantly different from 0. **(C-D)** Graphs show medians (bold lines), means (red diamonds), inner quartiles (boxes), and outer quartiles (whiskers) of thermal tolerance within each caste pooled across sites. Letters show the significance (different letters denotes p < 0.05) of pairwise differences in thermal tolerance among castes according to a post-hoc Tukey HSD test.

**Table 2.**
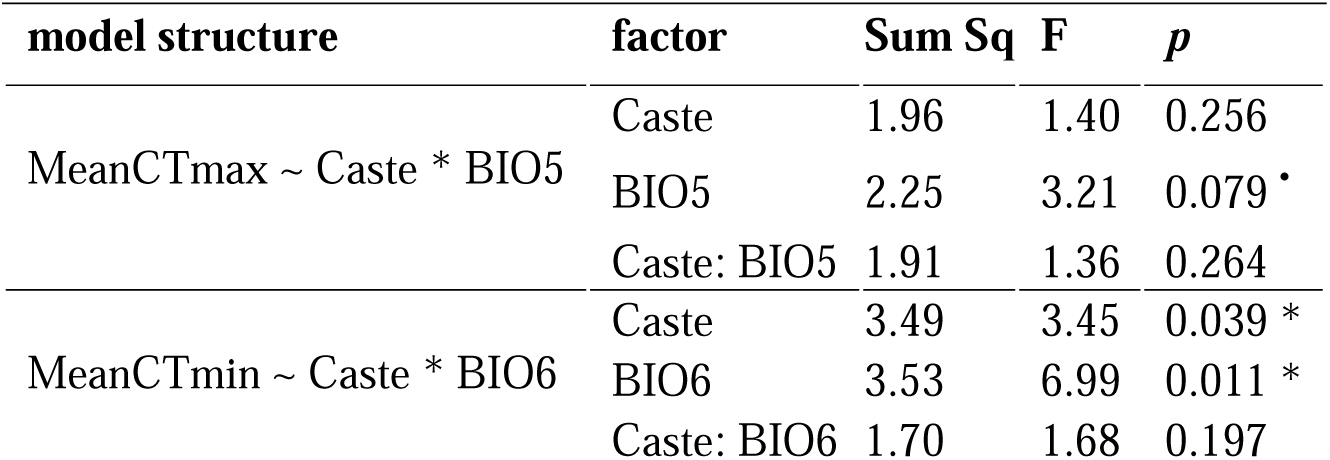
Statistical output of ANCOVA shown graphically in Figure 2. BIO5 is average highest temperature of warmest month. BIO6 is average lowest temperature of coldest month.

### Body size and lipid content as mechanistic explanators of cold tolerance

Wasp dry mass differed according to caste (F_2,54_ = 11.99, *p* < 0.001; Supplementary figure S1A) but showed no significant relationship to BIO6 (low temperature of the coldest month) (F_1,54_ = 0.47, *p* = 0.494). Results of a post-hoc Tukey HSD showed that foundresses were larger than workers (*p* < 0.001), though poorly sampled intermediates did not significantly differ in mass from foundresses (*p* = 0.372) or workers (*p* = 0.531). Among the assayed early-spring wasps, storage lipid mass was significantly lower for wasps living in colder climates (lower BIO6) (*X*^2^ = 4.45, df = 1, *p* = 0.035; Supplementary figure S1B), but storage lipid mass did not significantly differ across castes (*X*^2^ = 2.40, df = 2, *p* = 0.301).

However, CT_min_ was not a significant predictor of a wasp’s dry mass (F_1,54_ = 3.49, *p* = 0.067; Supplementary figure S1C) or the percentage of a wasp’s body mass composed of storage lipids (F_1,54_ = 1.46, *p* = 0.232).

### Heating tolerance & cooling tolerance across regions

The four surveyed regions in this study significantly differed in both warming tolerance (F_3,56_ = 9.00, *p* < 0.001) and cooling tolerance (F_3,54_ = 435.29, *p* < 0.001). Post-hoc Tukey HSD tests showed warming tolerance differences among regions that did not directionally follow latitude (Figure 3A; Supplementary Table S1) while cooling tolerance differences among regions were more highly significant and were also directionally predicted by latitude, with the northernmost site having some colonies with cooling tolerances below 0 (Figure 3B; Supplementary Table S2).

**Figure 3.**
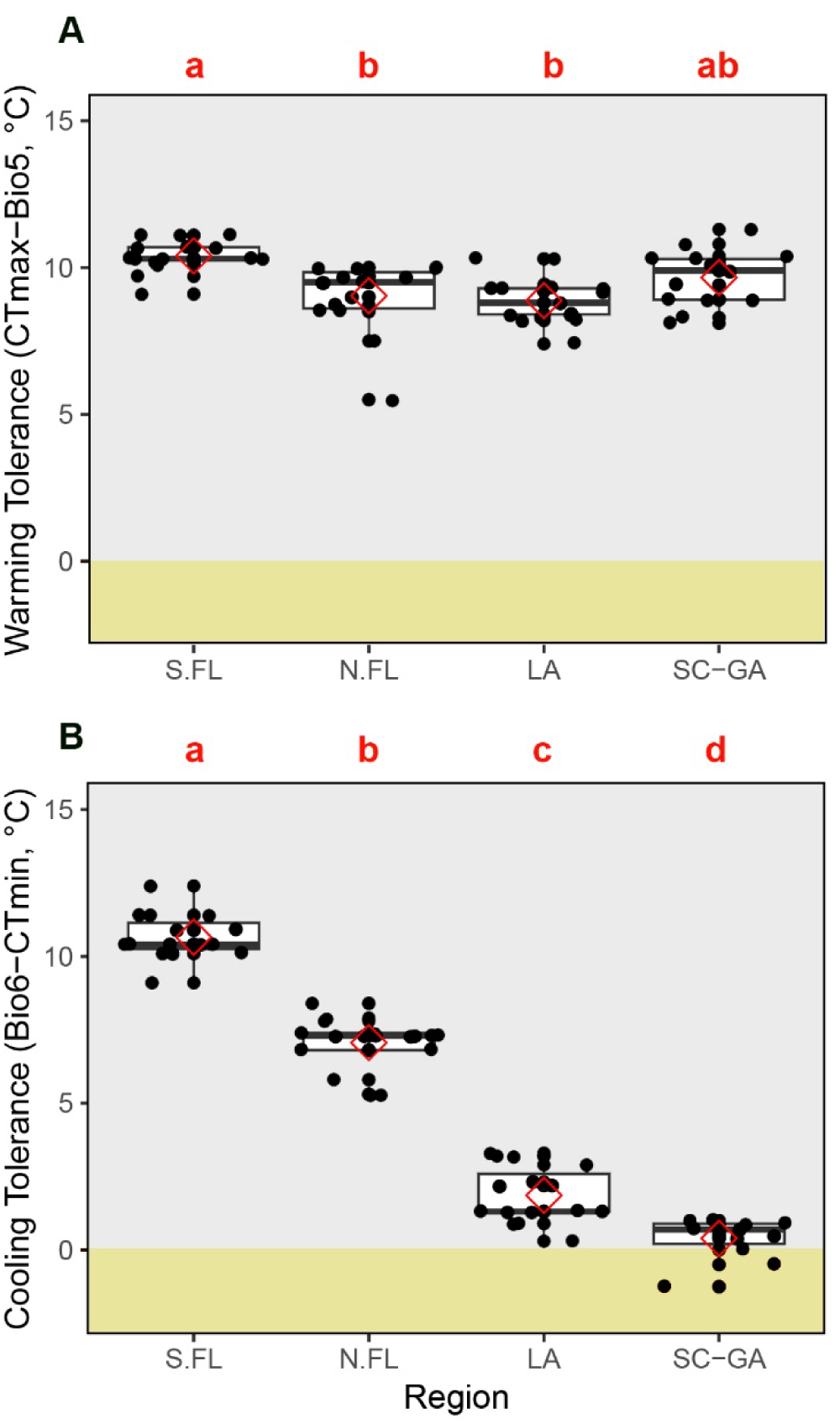
Differences in warming tolerance **(A)** and cooling tolerance **(B)** across regions. Regional median values are shown as bold lines, means are shown as red diamonds, inner quartiles are boxes, and outer quartiles are whiskers. Points represent averages for each caste within each colony. Letters show the significance (different letters denotes p < 0.05) of pairwise differences in thermal tolerance among castes according to a post-hoc Tukey HSD test.

## Discussion

### Geographic patterns

Although we report that cold tolerance (CT_min_) was significantly lower among wasps living in high latitude sites with more extreme low temperatures, heat tolerance (CT_max_) was not significantly predicted by environmental high temperatures, which may in part be due to there being less than 2° C variation in summertime high temperatures across the range of this mostly coastally distributed subspecies (Figure 1A). However, warming tolerances were also greater than 5° C across all sites (Figure 3A), suggesting that wasps in all regions are unlikely to be regularly experiencing mid-day heat coma under current and recent climatic conditions. Conversely, the proximity of lower critical temperatures to nighttime lows in the winter was well predicted by latitude, with northern wasps being very likely to experience nighttime cold coma in the winter (Figure 3B). This coupled with the predictive value of low temperatures on minimum critical temperature, is strong evidence that wintertime low temperatures are strong selection pressures on thermal performance.

For many eusocial insects, larger individuals tend to have higher CT_max_ and/or lower CT_min_ (Baudier et al., 2018; Baudier et al., 2015; Clemencet et al., 2010; Maebe et al., 2021; Wendt & Verble-Pearson, 2016), although this is not always the case (Kaspari et al., 2015; Oberg et al., 2012; Robinson & Baudier, 2024). *Mischocyttarus mexicanus cubicola* showed no difference in individual body size according to climate and wasp mass was not a significant predictor of CT_min_, suggesting that body size is not a main driver of the geographic patterns in cold tolerance (CT_min_) we report here. Although wasps from environments with lower winter temperatures did have relatively less lipid mass, there was no significant relationship between relative lipid mass and CT_min_. We therefore can conclude that these contrasts we observed in acute thermal performance are not likely to be driven by differences in fat stores, but that lipid stores play some other important role in overwintering for *M. mexicanus cubicula* colonies. Foundresses at higher latitudes may have had lower lipid body content in the spring because higher winter pressure caused them to metabolize a greater portion of their lipid stores and workers may similarly have emerged in early spring with lower fat stores. If this is indeed the case, then it would be consistent with patterns observed in several overwintering species of temperate *Polistes* (Stabentheiner et al., 2024; Yoshimura & Yamada, 2018). Disentangling the potential role of storage lipid content in the thermal adaptation of temperate and subtropical *Mischocyttarus* may thus necessitate future measurements across different seasons within each latitude.

### Caste & thermal performance

Caste was another important predictor of cold performance, with overwintering reproductive foundresses retaining mobility at lower temperatures than workers that emerged as adults from newly founded nests early in spring. Among eusocial insects, there can be group-level gains from dividing labor across task groups that specialize on different microclimates (Baudier & O’Donnell, 2017). For instance, some desert ants spread worker foraging effort across the day, with more heat tolerant ants foraging in midday while more sensitive individuals forage at less extreme times (Cerda & Retana, 1997). In the stingless bee *Tetragonisca angustula*, foragers that encounter more variable microclimates can also tolerate a wider range of temperatures than can standing or hovering guards that operate in the more thermally buffered microhabitat of the nest entrance (Robinson & Baudier, 2024). Cold-specialized reproductives have also been observed in other temperate social paper wasps, *Polistes exclamans* and *Polistes annularis*, with caste-associated differences in lower lethal temperatures and supercooling points (Strassmann et al., 1984). Here we report similar differences in CT_min_ for *M. mexicanus cubicola*.

Ontogenetic changes in thermal performance are common among holometabolous insects (Bowler & Terblanche, 2008). Callow adults being especially prone to cold coma has been observed in some ants (Baudier & O’Donnell, 2016), though there does not appear to be a relationship between whether an adult is callow and heat tolerance (Baudier & O’Donnell, 2016; Baudier et al., 2022). Because worker wasps collected in this study were younger than foundresses, we cannot rule out the possibility that freshly eclosed adult wasps might be more prone to cold coma than mature adults, regardless of caste. However, these potential age-based constraints do not preclude the adaptive value of these caste-specific differences. Caste specific differences in cold tolerance do not appear to be a consequence of body size or fat percentage, because during this sampling period foundresses and workers did not significantly differ in quantity of lipids stored, and although foundresses were larger in mass than workers there was no significant relationship between body size and CT_min_ (Supplementary Figure S1).

### Implications for climate change

In the context of climate change, cooling tolerance and warming tolerance results suggest that the thermal performance of *M. mexicanus cubicola* is more limited by low temperatures than by high temperatures across its studied range. Northward range expansion is therefore a possible consequence of local to global anthropogenic increases in temperature, while contraction at the southern end of this subspecies’ range is unlikely unless high temperatures increase more than 5° C. It should be noted, however, that this assessment does not include information about optimal performance temperatures, the thermal needs of developing brood, or the effects of less acute forms of thermal stress. The strategies used by high-latitude *M. mexicanus cubicula* to cope with more prolonged periods of cold coma are especially worth future exploration.

## Conclusion

The results of this study suggest that low temperature variation across the southeastern United States has shaped the thermal performance, and potentially the northward range expansion, of this social wasp.

Though this study presents evidence of this functional trait variation, a recent genotype by environment association study mirrors these patterns in this same system, showing strong predictive value of low temperature variation (Cook et al., 2023). Intriguingly, this work found two strong genetic clusters that correspond to the same northern and southern populations in this study, with an effect of climate, geography, and population structure towards local adaptation. Caste was also the strongest predictor for cold tolerance, which raises compelling questions about how individual functional traits interplay with cooperative nesting strategies across seasons in temperate versus subtropical populations. Together, this work fills important knowledge gaps between the fields of behavior and macrophysiology, justifying the use of flexible and disturbance-resilient model species in studies seeking to improve our understanding of the evolutionary selection pressures of climate and responses to anthropogenic climate change.

## Acknowledgements

This research was conducted using funds from the University of Rochester (to FMKU) and the University of Southern Mississippi (to KMR and KMB). We thank Alycia N. Johnson for assisting with lipid extractions. Permission to conduct research in Jean Lafitte Barataria Preserve was granted under permit number JELA-2021-SCI-0012. We thank the Friends of City Park for permitting collections made in New Orleans City Park. All other collections for this study were performed in public parking lots that required no permits.

## Author contributions

FMKU and KMB designed the study. KMB, FMKU, and KMR performed data collection, KMR and KMB conducted analyses, KMR and KMB drafted the manuscript, all authors contributed to manuscript revisions.

