## Supplementary Information for "Division of labor and low temperatures predict geographic variation in thermal tolerance of a North American paper wasp (*Mischocyttarus mexicanus cubicola*)"

**Table S1.** Output of post-hoc Tukey HSD test conducted to compare warming tolerance among the four regions studied. For a graph of these results see Figure 3A.

| **Sites** | **Estimate** | **Std. Error** | ***t*** | ***p*** |
| --- | --- | --- | --- | --- |
| N.FL – LA | 0.1635 | 0.3213 | 0.509 | 0.9566 |
| S.FL – LA | 1.5235 | 0.3213 | 4.741 | <0.001 * |
| SC-GA – LA | 0.7851 | 0.3342 | 2.349 | 0.0991 . |
| S.FL – N.FL m | 1.3600 | 0.3312 | 4.106 | <0.001 * |
| SC-GA – N.FL | 0.6215 | 0.3437 | 1.808 | 0.2800 |
| SC-GA – S.FL | -0.7385 | 0.3437 | -2.148 | 0.1505 |

**Table S2.** Output of post-hoc Tukey HSD test conducted to compare cooling tolerance among the four regions studied. For a graph of these results see Figure 3B.

| **Sites** | **Estimate** | **Std. Error** | ***t*** | ***p*** |
| --- | --- | --- | --- | --- |
| N.FL – LA | 5.1988 | 0.2980 | 17.445 | <0.001 * |
| S.FL – LA | 8.7933 | 0.3072 | 28.626 | <0.001 * |
| SC-GA – LA | -1.4645 | 0.3339 | -4.386 | <0.001 * |
| S.FL – N.FL m | 3.5945 | 0.2980 | 12.062 | <0.001 * |
| SC-GA – N.FL | -6.6634 | 0.3255 | -20.469 | <0.001 * |
| SC-GA – S.FL | -10.2579 | 0.3339 | -30.717 | <0.001 * |

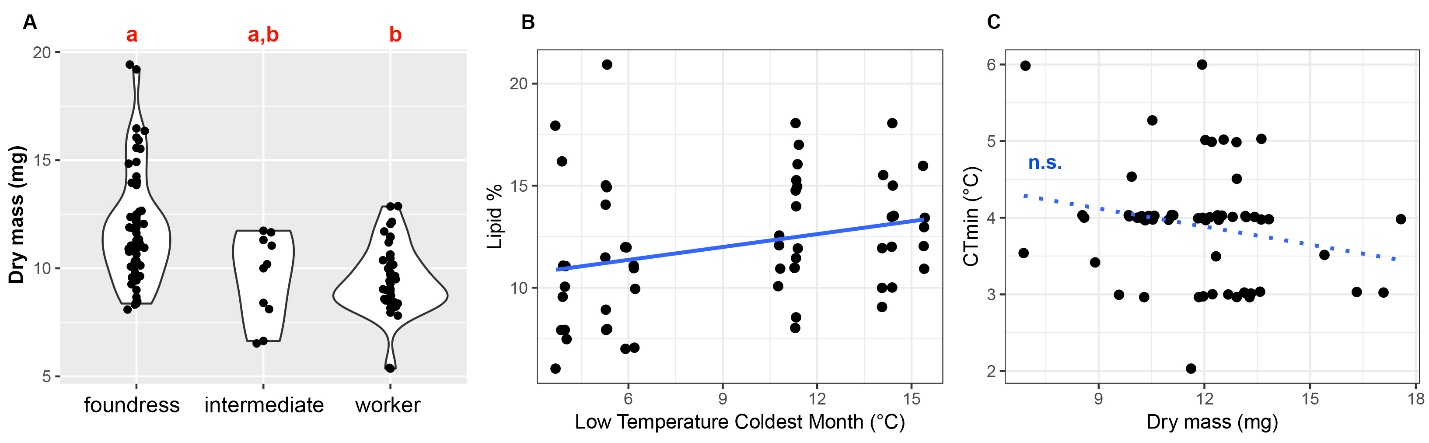

**Figure S1.** Graphs exploring body size and percent storage lipids as possible explanators of the climatically-driven patterns in CT_min_. Statistical outputs are reported in the results section of the main paper. **(A)** Significantly greater dry mass of foundresses relative to workers **(B)** Significantly lower body proportional fat stores in more low-temperature experiencing wasps. **(C)** Marginally non-significant relationship between body size and CT_min_.
